# Direct Cell Extraction of Membrane Proteins for Structure-Function Analysis

**DOI:** 10.1101/2022.07.05.498330

**Authors:** Ieva Drulyte, Aspen Rene Gutgsell, Pilar Lloris-Garcerá, Michael Liss, Stefan Geschwindner, Mazdak Radjainia, Jens Frauenfeld, Robin Löving

## Abstract

Membrane proteins are the largest group of therapeutic targets in a variety of disease areas and yet, they remain particularly difficult to investigate. We have developed a novel one-step approach for the incorporation of membrane proteins directly from cells into lipid Salipro nanoparticles. Here, with the pannexin1 channel as a case study, we demonstrate the applicability of this method for structurefunction analysis using SPR and cryo-EM.

## Main

Membrane proteins are central to many physiological processes and are of critical importance for the pharmaceutical industry. More than 60% of all current clinically-approved drugs target proteins embedded in the lipid membrane^1,2^. However, membrane proteins are particularly difficult to investigate in drug discovery campaigns because they are notoriously difficult to purify using standard purification methods.

To address this problem, the Salipro nano-membrane system was developed as a universal platform to stabilize membrane proteins in a native-like lipid environment using a scaffold of saposin proteins^3–6^. The Salipro platform has recently expanded to enable the direct extraction of sensitive membrane proteins from crude cell membranes, thereby eliminating the need for detergent-purification^7^. This methodology is termed DirectMX for direct membrane extraction. In this work, we present a case study using a streamlined version of the DirectMX protocol working directly from crude cell pellets to purify the ATP release channel pannexin1 (PANX1), and investigate its structural and ligand-binding properties.

PANX1 is a potential therapeutic target and associated with various pathologies, including inflammation, pain, ischemia, and epilepsy^8,9^. A major limitation for exploring PANX1 biology, or its potential as a therapeutic target, is an overall lack of biophysical studies using isolated systems. To date, several pharmacological agents have been reported to inhibit PANX1 using cellular functional assays; however, it is impossible to distinguish if the observed phenotypic effects are directly from PANX1 target engagement or due to indirect effects from binding to other proteins. Here, we aimed to use the Salipro DirectMX method, working directly from cell pellets, to isolate PANX1 and perform the first-ever *in vitro* structurefunction assays using surface plasmon resonance (SPR) and single-particle cryoelectron microscopy (cryo-EM).

To begin, expression constructs for mouse PANX1 (mPANX1) containing cleavable C-terminal GFP and TwinStep affinity tags were transiently expressed in Expi293 cells, harvested 48 hours post-transfection, and stored at −80°C. Cell pellets were resuspended in digitonin-containing buffer to increase membrane fluidity and render lipids and membrane proteins more accessible for reconstitution using saposin A (Figure 1A). Following a 10-minute incubation with saposin A, the formation of saposin-containing mPANX1-GFP particles was assessed by analytical size exclusion chromatography (SEC) using a fluorescens detector (Figure 1B). For affinity purification, affinity-tagged mPANX1-containing particles were immobilised on a StrepTactin Sepharose column and eluted using the PreScission protease for on-column cleavage (Figure 1C). Next, the eluate was concentrated by ultrafiltration and subjected to preparative SEC (Figure 1D). SEC fractions were analyzed by silver staining after SDS-PAGE revealed the formation of pure and homogenous Salipro-mPANX1 nanoparticles (Figure 1E). SEC fractions containing Salipro-mPANX1 were pooled, concentrated and flash frozen prior to use in downstream applications.

**Figure 1.**
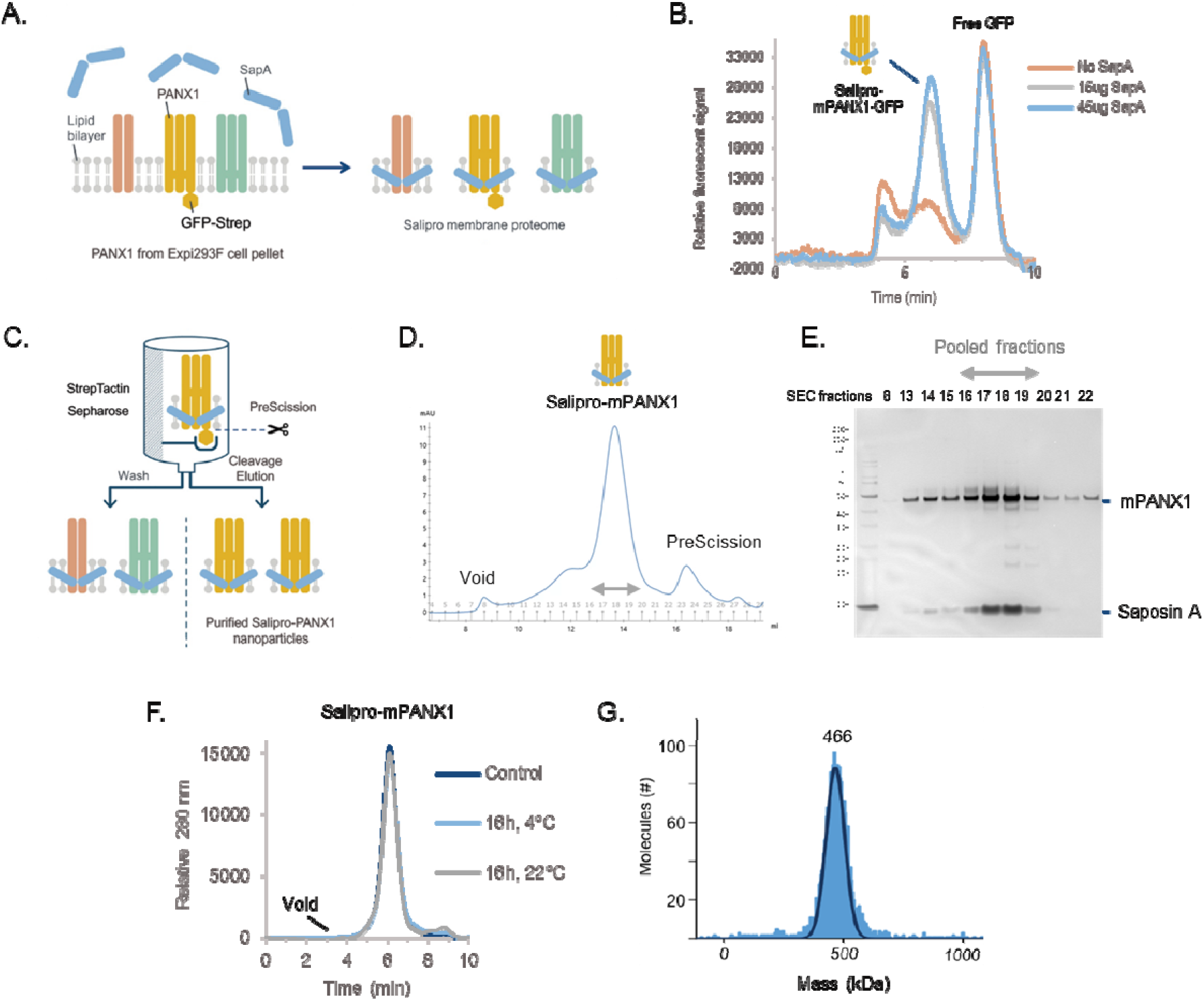
Salipro-mPANX1 DirectMX summary. (A) Schematic illustration of the DirectMX methodology. (B) mPANX1 expressed as a GFP fusion protein formed a homogenous population of Salipro-mPANX1-GFP nanoparticles upon addition of SapA, peaking around 6 min. (C) Schematic illustration of the Salipro-mPANX1 affinity purification using StrepTactin Sepharose with on-column PreScission cleavage. (D) Salipro-mPANX1 particle purification by preparative SEC. (E) SEC fraction analysis by SDS-PAGE. (F) Analytic SEC trace of Salipro-mPANX1 particles that were thawed and incubated for 16h at 4°C or 22°C. The control sample was analyzed immediately after freeze-thawing without any further incubation. (G) Mass photometry confirms the presence of a homogenous population of Salipro-mPANX1 particles with an apparent nanoparticle size of 466 kDa.

Structure-function analysis of membrane proteins require a stable sample and we therefore assessed the thermal stability of freeze-thawed Salipro-mPANX1 particles at 4°C or 22°C for 16 hours using analytic SEC (Figure 1F). Salipro-mPANX1 particles remained stable and homogenous at 22°C for up to 16 hours (Figure 1F). Mass photometry measurements^10^ also showed a homogenous population of freeze-thawed Salipro-mPANX1 particles at a molecular size of approximately 466 kDa (Figure 1G). Analogous to the DirectMX extraction of mPANX1, we applied the same methodology for human PANX1 (hPANX1) expressed in sf9 insect cells, leading to similar results (Supplementary figure 1).

We then employed SPR to directly assess the ligand-binding competence of purified Salipro-PANX1 particles. In the case of membrane proteins, SPR analysis remains challenging due to the intrinsic instability of native membrane proteins solubilized in detergents and the difficulty to immobilize enough functional membrane proteins to obtain significant binding signal intensity when working with small molecular ligands^11^. Untagged mPANX1 embedded in a His-tagged Salipro nanoparticle was immobilized (Figure 2A) and challenged with benzoylbenzoyl-ATP (bzATP), spironolactone, and carbenoxolone. Binding constants were determined for bzATP (K_D_ 720 μM ± 133 μM) and spironolactone (K_D_ 160 μM ± 10 μM) (Figure 2B-C). Binding to carbenoxolone was also confirmed, although the binding constant could not be determined (Supplementary figure 2). Similar results were observed for biotin-Salipro-hPANX1 nanoparticles (Supplementary figure 3) with all SPR data summarized in Supplementary table 1. In summary, the downstream analysis showed a high degree of functional Salipro-PANX1 material immobilized on the sensor surface with an approximate binding capacity to small molecular compounds between 80-90%.

**Figure 2.**
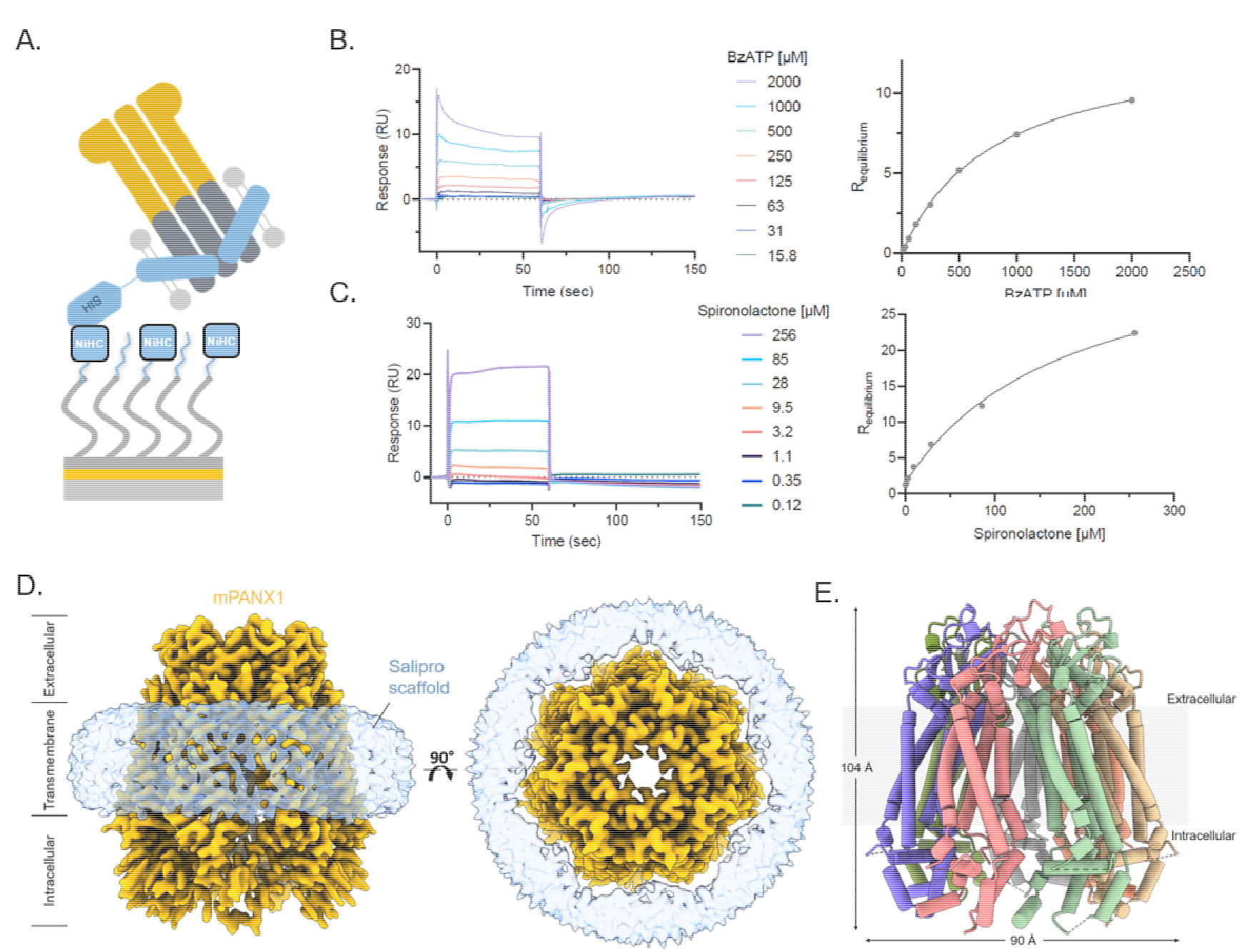
Salipro-mPANX1 structure and functional analysis. (A) Schematic of His-Salipro-mPANX1 nanoparticles immobilisation to a sensor with Ni^2+^ ions (NiHC) binding to the His-tag. Concentration series of bzATP (B) and spironolactone (C) was injected over a His-Salipro-mPANX1 coated surface with a calculated KD of 720 μM ± 133 μM and 160 μM ± 10 μM, respectively. (D) Cryo-EM reconstruction of Salipro-mPANX1 at 3.1Å. The ion channel (orange) and Salipro lipid disk (blue) is shown as side view (left) and top view (right). (E) The structural model of mPANX1 with each subunit colored. All SPR data is representative of n=3.

Lastly, we present the first cryo-EM structure of a membrane protein that was directly reconstituted from crude cell pellets into Salipro nanoparticles (Figure 2D-E). To date, all cryo-EM structures of membrane proteins in Salipro particles were obtained from membrane proteins that were detergent-purified first and, in a second step, reconstituted with an artificial mix of lipids^3–6^. Briefly, cryo-EM grids were prepared using Salipro-mPANX1 nanoparticles and standard Vitrobot plunge-freezing. Cryo-EM micrographs revealed that frozen-hydrated complexes were monodisperse with 2D class averages diplaying secondary and tertiary structures, as well as heptameric organization of mPANX1 (Supplementary figure 4). Using a total of 269,000 particles, the 3D structure reconstruction reached a 3.1 Å resolution (Supplementary Figure 5and Supplementary table 2).

Extracellular and transmembrane regions of the Salipro-mPANX1 are well-resolved in the structure with clear side-chain density visible (Supplementary Figure 6A). In contrast, much of the intracellular helices were not well resolved in the cryo-EM map prohibiting complete atomic modelling. We next compared Salipro-mPANX1 to hPANX1 (PDB code 6WBF15), a homologue with a shared sequence identify of 86.2 %. The dimensions of the Salipro-mPANX1 heptamer are 90 by 104 Å (Figure 2E), which is slightly smaller than those of hPANX1, which are 96 by 109 Å. On the protomer level, the mouse and human PANX1 structures are highly similar, with an root-mean-square deviation (RMSD) of 1.556 across 256 aligned atom pairs. The intracellular region of the Salipro-mPANX1 exhibits the most differences compared to the hPANX1 with several alpha-helices shifted; for example, helices encompassing residues 342-349 (Salipro-mPANX1) and 342-355 (hPANX1) reach a maximum displacement of ~5 Å (Supplementary Figures 6B and 6C).

In this proof-of-concept study, we present the Salipro method for the reconstitution and purification of homogenous wildtype mPANX1 and hPANX1 particles directly from cell pellets. With this, the first PANX1 *in vitro* small molecular ligand-binding assay was developed, enabling screening compounds for this drug target via SPR. The cryo-EM structure of mouse PANX1 was solved (3.1 Å) for the first time, also representing the first published structure of a membrane protein extracted by Salipro nanoparticles directly from cells. This lays the groundwork for future studies to investigate novel PANX1-binders on a structural and functional level by SPR and cryoEM.

The direct cell extraction presented here enables purification of stable and functional wildtype membrane proteins without the need for laborious and time-consuming detergent screenings, protein engineering or screening of applicable alternative scaffolding setups. We believe that the Salipro direct cell extraction may accelerate future membrane protein research, as well as enable the development of new therapeutic ligands in a time-efficient and streamlined manner.

## Online methods

### Material and Reagents

Human saposin A was recombinantly produced and purified as previously described ^1,2^. Pure saposin A in HNG buffer (50 mM HEPES at pH 7.5, 150 mM NaCl, 5% (v/v) glycerol) was stored at −80°C. Saposin A empty nanoparticles (Salipro-Empty particles) were prepared as previously described^1^. PreScission protease was purchased from Cytiva. PANX1 ligands for SPR binding measurements were Benzoylbenzoyl-ATP (Sigma-Aldrich, CAS 112898-15-4), Carbenoxolone (sourced at AstraZeneca) and Spironolactone (sourced at AstraZeneca).

### PANX1 expression

Two PANX1 expression constructs were designed to express mPANX1 and hPANX1. The mPANX1 construct was designed to contain the wildtype mouse PANX1 sequence with a n-terminal FLAG tag and a c-terminal GFP fused to a Twin Strep tag. In addition, the mPANX1 sequence was designed to be flanked by two PreScission protease cleavage sites (FLAG-3C-mPANX1-3C-GFP-StrepII) enabling purification of mPANX1 without the presence of GFP and tags in the final preparation. The expression construct was codon optimised for protein expression in human cells. Human Expi293 cells were transiently transfected, and cell pellets were collected 5 days post transfection (GeneArt, Thermo Fisher Scientific). The hPANX1 construct was designed to contain the wildtype human PANX1 sequence with a c-terminal eGFP fused to a His^10^ tag followed by an EPEA affinity tag. In addition, a PreScission protease cleavage site was designed directly downstream of hPANX1 (hPANX1-3C-GFP-His^10^-EPEA) enabling purification of hPANX1 without the presence of GFP and tags in the final preparation. The expression construct was codon optimised for protein expression in insect cells. Viral stocks were prepared and used to induce protein expression in sf9 insect cells. Cell pellets of hPANX1 expressing cells were collected 48 hours post infection (GeneArt, Thermo Fisher Scientific). All cell pellets expressing PANX1 were frozen and stored at −80 °C.

### Membrane protein reconstitution screening using fluorescence-detection size-exclusion chromatography (FSEC)

Cell pellet expressing mPANX1-GFP was thawed and resuspended with a five-time excess of HNG buffer supplemented with 1.2x cOmplete protease inhibitor cocktail (Roche) and 1.2% digitonin (Calbiochem) to increase membrane fluidity^3^. All steps were performed at 4 °C. The mixture was incubated for one hour in a rotating wheel, followed by centrifugation at 30,000 g for 45 minutes to remove insoluble material. Saposin A (15 or 45 μg) was added to the lysate mixture in a total volume of 50ul, followed by a 10 min incubation. One sample lacking Saposin A served as negative control. The formation of saposin A reconstituted mPANX-GFP nanoparticles (Salipro-mPANX1-GFP) was evaluated by analytic size exclusion chromatography (SEC) using a fluorescent detector (FSEC) in a Superose 6 Increase 5/150 GL column (Cytiva) equilibrated with detergent free HNG buffer, using a Prominence-i LC-2030C high performance liquid chromatography system equipped with PDA and RF-20Axs fluorescence detectors (Shimadzu) at a flow rate of 0.3 mL/min. In-line light absorbances were monitored for total protein elution at 280 nm and for GFP elution at 512 nm, with excitation for the latter set to 500 nm.

### Salipro-PANX1 particle purification and characterization

For affinity chromatography to purify Salipro-mPANX1, 500 μL of Strep Tactin Sepharose High Performance resin (GE Healthcare) were equilibrated with HNG buffer in a Poly-prep® Chromatography column (Bio-Rad). The lysate mixture described above was loaded to the column and the sample flow-through was passed 3 more times over the resin. The PANX1-loaded affinity resin was incubated at 4 °C with 45 mg of Saposin A and after 30 min the flow-through was discarded. After washing the affinity resin with HNG buffer, the beads with bound Salipro-PANX1-GFP were resuspended with HNG buffer containing 60U of PreScission protease and incubated at 4 °C overnight in a rotating wheel. After cleavage, the mixture was loaded back into the empty Poly-prep® column and purified Salipro-mPANX1 particles were eluted in the flow-through, while the GFP-tag remained bound to the resin. The elution was concentrated with Amicon filter device with 50-kDa NMWL (Millipore) and subjected to preparative SEC in a Superose 6 Increase 10/300 GL column (Cytiva), equilibrated in HNG buffer, using an Äkta Pure chromatography system (Cytiva) at a flow rate of 0.35 mL/min. The fractions were collected and analysed by SDS-PAGE using NuPAGE 4-12% Bis-Tris polyacrylamide gels (Thermo Fisher Scientific) and visualized by Silver Stain. Fractions containing pure Salipro-mPANX1 nanoparticles were pooled, concentrated as described above, flash frozen and stored at −80 °C. Salipro-hPANX1 particles were made using a similar protocol were the hPANX1 expressing sf9 cells were treated with 1% DDM to increase membrane fluidity prior to Salipro reconstitution followed by affinity chromatography using the CaptureSelect™ C-tag Affinity Matrix (Thermo Fisher Scientific).

To evaluate particle stability, purified Salipro-mPANX1 and Salipro-hPANX1 particles were thawed after flash freezing and incubated for 16 hours at either 4 °C or 22 °C prior to analytic SEC.

### Saposin A labelled Salipro-PANX1 particles

To generate His-tagged or biotinylated Salipro-PANX1 particles, the same method as described above was used to extract and purify Salipro-PANX1 using His-tagged or biotinylated Saposin A. His-tagged Saposin A was produced and purified as previously described^1,2^, albeit without the TEV protease cleavage step for His-tag removal. Saposin A was biotinylated in a one-hour incubation step at RT using NHS-PEG12-Biotin (Thermo Fisher Scientific), following the manufacturer’s recommendations. The Quant*Tag™ Biotin Kit (Vector Laboratories) was used to confirm the amount of saposin A biotinylation.

### SPR analysis

All SPR experiments were conducted on a BIAcore S200 or T200 instrument (Cytiva) using 10 mM HEPES pH 7.4, 150 mM NaCl (HBS-N) as the running buffer.

#### Salipro-mPANX1 SPR Assays

Multicycle kinetic experiments were conducted using a sensor with Ni^2^+ ions complexed on a two-dimensional chelating surface (NiHC 1500M, Xantec Bioanalytics GmbH) at 12°C. Prior to ligand immobilization, the sensor was washed with 300 mM EDTA pH 8.3 and loaded with Ni^2^+-ions by injecting a 500 nM solution of NiCl2 in running buffer for 3 min. His-tagged saposin A empty nanoparticles (His-Salipro-empty) and his-tagged Saposin A mPANX1 particles (His-Salipro-mPANX1) were injected at a concentration of 10-20 ng/mL using a contact time of 16 min at 1μL/min to achieve desired densities of 2500-4000 RU. His-Salipro-Empty were immobilized on the reference surface as a lipid-only control. Increasing concentrations of (0.12, 0.35, 1.1, 3.2, 9.5, 28, 85, and 265 μM) of carbenoxolone or spironolactone or (0.03, 0.06, 0.13, 0.25, 0.50, 1.00, 2.00, 4.00 mM) of benzoylbenzoyl-ATP were iteratively injected with a 60 s contact time, followed by a 90 s dissociation phase. All resulting sensorgrams were reference and blank subtracted prior to fitting.

#### Biotin-Salipro-hPANX1 SPR Assays

Multicycle kinetic experiments were conducted using a biotin-coated sensor (Cytiva) at 12°C. Prior to ligand immobilization, the sensor was washed with 50 mM NaOH, 1 M NaCl for 1 min. Biotin-tagged saposin A empty nanoparticles (Biotin-Salipro-Empty) and biotin-tagged saposin A hPANX1 particles (Biotin-Salipro-hPANX1) were injected at a concentration of 10-20 ng/mL using a contact time of 16 min at 1μL/min to achieve desired densities of 2500-4000 RU. Biotin-Salipro-Empty particles were immobilized on the reference surface as a lipid-only control. Increasing concentrations of (2, 4, 8, 16, 32, 64, 128, and 265 μM) of carbenoxolone or spironolactone or (0.03, 0.06, 0.13, 0.25, 0.50, 1.00, 2.00, 4.00 mM) of benzoylbenzoyl-ATP were iteratively injected with a 60 s contact time, followed by a 90 s dissociation phase. All resulting sensorgrams were reference and blank subtracted prior to fitting.

### Mass Photometry

All samples were measured in 1X HBS-N buffer (10 mM HEPES pH 7.4, 150 mM NaCl) using the Refeyn OneMP mass photometer (Refeyn Ltd.) with a 60 s acquisition time. The resulting histograms were fitted to Gaussian distributions using DiscoverMP (Refeyn Ltd.) to extract peak contrast and relative amount of each peak (n=3). Contrast-to-mass conversion was achieved by calibration using NativeMark protein ladder (Thermo Fisher Scientific). Three protein species (with specified masses) were fitted to corresponding Gaussian distributions to extract a linear relation between mass and contrast.

### Cryo-EM sample preparation and data collection

Quantifoil 200 mesh 1.2/1/3 Cu grids were glow-discharged using 20 mAmp current for 45 sec and charge set to positive (GloQube®, Quorum Technologies). To overcome preferred orientation, a purified full-length Salipro-mPANX1 protein sample was mixed with 0.5 mM Fos-Choline 8 (Anatrace) before the grid freezing. 3 ul of 2.5 mg/ml protein was pipetted onto the grid in Vitrobot Mark IV (Thermo Fisher Scientific) chamber set to 4 °C and 95% humidity. Grids were then blotted for 10 s using a blot force of +20 and 30 s waiting time before plunge-freezing in liquid ethane.

Data collection was conducted using a 300 kV Thermo Scientific™ Krios G4™ Cryo-Transmission Electron Microscope (Cryo-TEM) equipped with Selectris X Imaging Filter and Falcon 4™ Direct Electron Detector camera operated in Electron-Event representation (EER) mode. Thermo Fisher Scientific EPU 2 software was used to automate the data collection. Aberration-free image shifts (AFIS) and Fringe-free Imaging (FFI) increased the data collection throughput to ~360 images/hour. 7995 movie dataset was collected in EFTEM mode using 10 eV slit and a nominal magnification of 165,000x, resulting in a calibrated pixel size of 0.727 Å/px. The exposure time of 4.16 s with a total dose of 40.24 e^-^ /Å^2^ was used, and each movie was split into 40 fractions during motion correction. The dose rate on the camera was 5.4 e^-^/px/s, and the nominal defocus range was specified between −0.5 and −1.5 μm in 0.25 μm intervals.

### Cryo-EM single-particle analysis

Data processing was performed using Relion 3^4^ and CryoSPARC™ ^5^ image processing suites. After motion- and CTF-correction (using Relion’s implementation of MOTIONCOR^6^ and CTFFIND-4.1^7^, respectively), particles were picked using a template-free auto-picking procedure based on a Laplacian-of-Gaussian (LoG) filter with particle diameter set to 130-170 Å. A total of 1,314,251 particles were picked from 6739 micrographs and exacted in a 320 px box Fourier cropped 4x to 80 px box. The initial model was generated in cryoSPARC™, performing the initial pre-processing steps described above independently to the Relion workflow. Following one round of 2D and 3D classification (the latter with C1 symmetry), the best 3D class, displaying clear secondary structure features and consisting of 307,098 particles, was chosen. Refinement using C7 symmetry and solvent flattening resulted in a 3.4 Å reconstruction. This stack of particles was subjected to CTF refinements, Bayesian polishing and further two 3D classification rounds without particle alignment, after which the best 268,823 particles were selected. Subsequent refinement, masking and sharpening yielded the final 3.1 Å map. The map resolution was determined based on the gold-standard 0.143 criterion.

### Model building, validation, and structural analysis

AlphaFold structure prediction of Mus musculus Pannexin-1 (AF-Q9JIP4-F1-model_v2.pdb) was used as a starting model^8,9^. Model re-building was performed in Coot^10^ and ISOLDE^11^. The final model was refined with anisotropic atomic B-factors in reciprocal space using REFMAC^12^. Molprobity score, clash score, rotamer and Ramachandran analyses were performed using MOLPROBITY^13^.

## References

1. Overington, J. P., Al-Lazikani, B. & Hopkins, A. L. How many drug targets are there? Nat. Rev. Drug Discov. 5, 993–996 (2006).

2. Rask-Andersen, M., Almén, M. S. & Schiöth, H. B. Trends in the exploitation of novel drug targets. Nat. Rev. Drug Discov. 10, 579–590 (2011).

3. Frauenfeld, J. et al. A saposin-lipoprotein nanoparticle system for membrane proteins. Nat. Methods 13, 345–351 (2016).

4. Nguyen, N. X. et al. Cryo-EM structure of a fungal mitochondrial calcium uniporter. Nature 559, 570–574 (2018).

5. Kim, J. J. et al. Shared structural mechanisms of general anaesthetics and benzodiazepines. Nature 585, 303–308 (2020).

6. Lyons, J. A., Bøggild, A., Nissen, P. & Frauenfeld, J. Saposin-Lipoprotein Scaffolds for Structure Determination of Membrane Transporters. in Methods in Enzymology vol. 594 85–99 (Elsevier, 2017).

7. Lloris-Garcerá, P. et al. DirectMX – One-Step Reconstitution of Membrane Proteins From Crude Cell Membranes Into Salipro Nanoparticles. Front. Bioeng. Biotechnol. 8, 215 (2020).

8. Whyte-Fagundes, P. & Zoidl, G. Mechanisms of pannexin1 channel gating and regulation. Biochim. Biophys. Acta BBA - Biomembr. 1860, 65–71 (2018).

9. Sanchez-Arias, J. C. et al. Purinergic signaling in nervous system health and disease: Focus on pannexin 1. Pharmacol. Ther. 225, 107840 (2021).

10. Sonn-Segev, A. et al. Quantifying the heterogeneity of macromolecular machines by mass photometry. Nat. Commun. 11, 1772 (2020).

11. Geschwindner, S., Carlsson, J. F. & Knecht, W. Application of Optical Biosensors in Small-Molecule Screening Activities. Sensors 12, 4311–4323 (2012).

## Online Method References

1. Frauenfeld, J. et al. A saposin-lipoprotein nanoparticle system for membrane proteins. Nat Methods 13, 345–351 (2016).

2. Lyons, J. A., Bøggild, A., Nissen, P. & Frauenfeld, J. Saposin-Lipoprotein Scaffolds for Structure Determination of Membrane Transporters. in Methods in Enzymology vol. 594 85–99 (Elsevier, 2017).

3. Lloris-Garcerá, P. et al. DirectMX – One-Step Reconstitution of Membrane Proteins From Crude Cell Membranes Into Salipro Nanoparticles. Front. Bioeng. Biotechnol. 8, 215 (2020).

4. Scheres, S. H. W. RELION: Implementation of a Bayesian approach to cryo-EM structure determination. Journal of Structural Biology 180, 519–530 (2012).

5. Punjani, A., Rubinstein, J. L., Fleet, D. J. & Brubaker, M. A. cryoSPARC: algorithms for rapid unsupervised cryo-EM structure determination. Nat Methods 14, 290–296 (2017).

6. Zheng, S. Q. et al. MotionCor2: anisotropic correction of beam-induced motion for improved cryo-electron microscopy. Nat Methods 14, 331–332 (2017).

7. Rohou, A. & Grigorieff, N. CTFFIND4: Fast and accurate defocus estimation from electron micrographs. Journal of Structural Biology 192, 216–221 (2015).

8. Tunyasuvunakool, K. et al. Highly accurate protein structure prediction for the human proteome. Nature 596, 590–596 (2021).

9. Varadi, M. et al. AlphaFold Protein Structure Database: massively expanding the structural coverage of protein-sequence space with high-accuracy models. Nucleic Acids Res 50, D439–D444 (2022).

10. Emsley, P. & Cowtan, K. Coot: model-building tools for molecular graphics. Acta Cryst D 60, 2126–2132 (2004).

11. Croll, T. I. ISOLDE: a physically realistic environment for model building into low-resolution electron-density maps. Acta Cryst D 74, 519–530 (2018).

12. Murshudov, G. N. et al. REFMAC5 for the refinement of macromolecular crystal structures. Acta Cryst D 67, 355–367 (2011).

13. Williams, C. J. et al. MolProbity: More and better reference data for improved all-atom structure validation. Protein Science 27, 293–315 (2018).

