## Supplementary data for "Direct Cell Extraction of Membrane Proteins for Structure-Function Analysis"

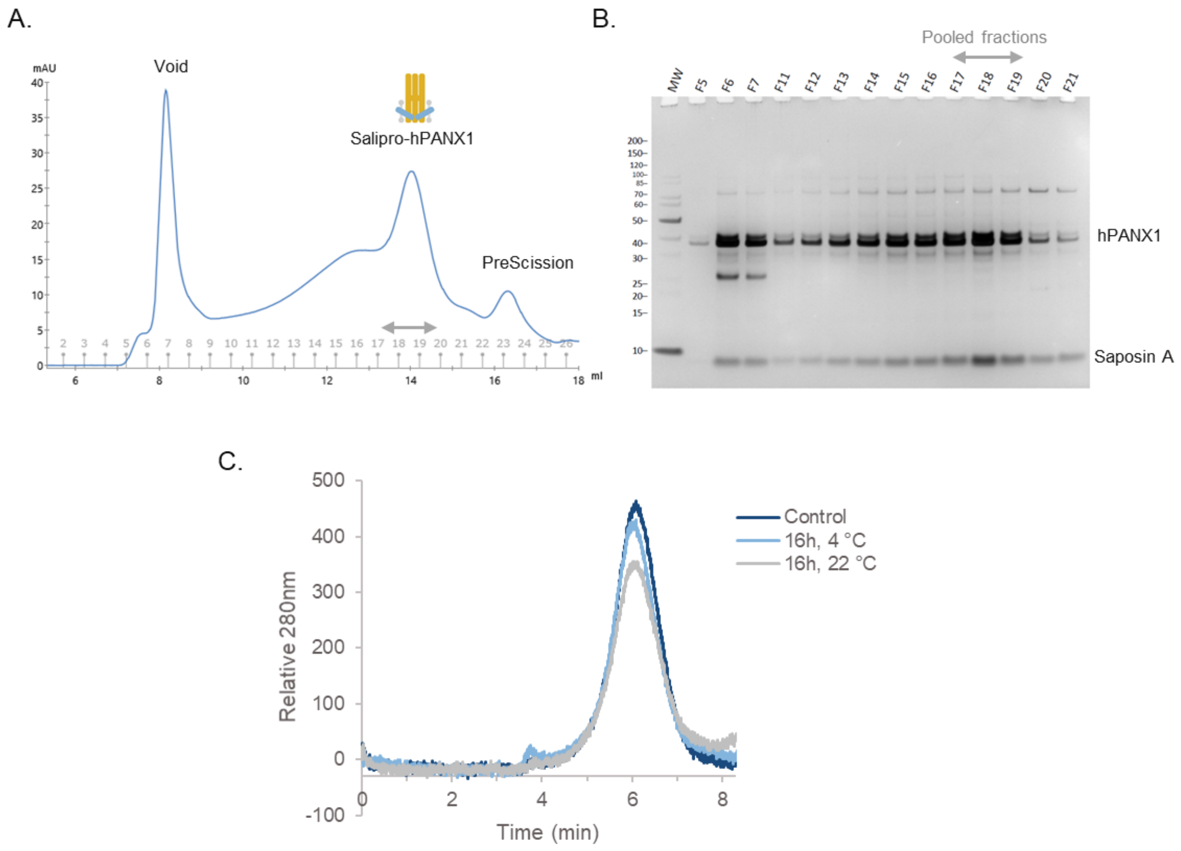

**Supplementary figure 1.** Direct cell extraction of Salipro-hPANX1. The hPANX1-GFP fusion protein was designed to contain c-terminal $^{10}$His/EPEA affinity tags with a PreScission protease cleavage site upstream of GFP (hPANX1-PreScission-GFP-$^{10}$His/EPEA). Protein purification was achieved using C-tag affinity purification resin binding to the EPEA tag, followed by on-column PreScission cleavage. (A) Eluted Salipro-hPANX1 particles were concentrated by ultrafiltration and further purified by preparative SEC. (B) Collected SEC fractions were analysed by reducing SDS-PAGE using protein silver staining confirming the sole presence of hPANX1 and saposin A making up the Salipro-hPANX1 nanoparticles. SEC peak fractions 17-20 were pooled, concentrated and flash frozen in liquid nitrogen for -80 °C storage. (C) To evaluate Salipro-hPANX1 stability, particles were thawed and incubated for either 16h at 4 °C or at 22 °C followed by analytic SEC. The control sample was analyzed immediately after freeze-thawing without any further incubation.

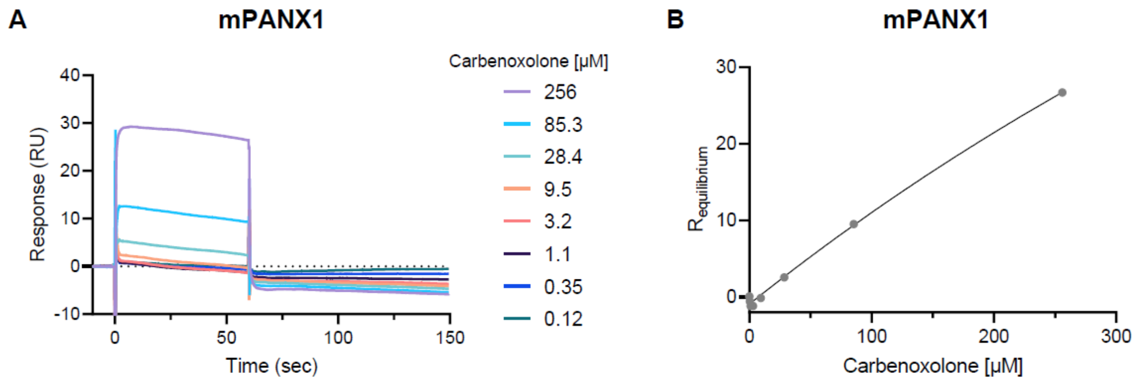

**Supplementary figure 2.** SPR analysis of His-Salipro-mPANX1 particle binding to carbenoxolone (A) Concentration series of carbenoxolone was injected over a His-Salipro-mPANX1 coated surface for 60 seconds followed by a 90 second dissociation time. (B) The equilibrium curve does not have sufficient curvature to accurately determine the KD. Data is representative of n=3.

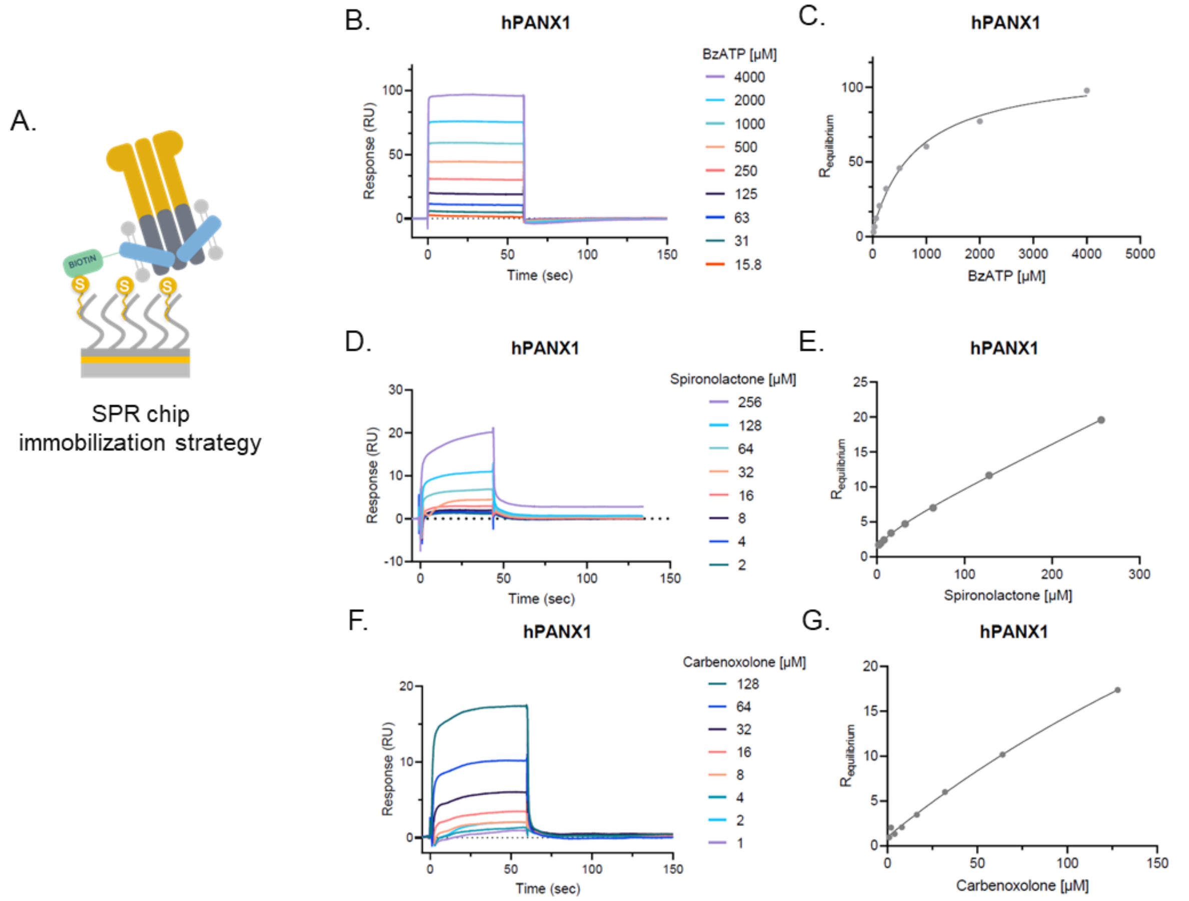

**Supplementary figure 3.** SPR analysis using biotin-Salipro-hPANX1 particle (A) Schematic illustration of biotin-Salipro-hPANX1 nanoparticles immobilization to a sensor chip coated with streptavidin (S). (B) Concentration series of BzATP was injected over a human PANX1 coated surface for 60 seconds followed by a 90 second dissociation time. (C) Equilibrium analysis showed a KD of 837µM ± 140µM. (D) Concentration series of spironolactone was injected over the surface for 45 seconds followed by a 90 second dissociation time. (E) The equilibrium curve does not have sufficient curvature to accurately determine the KD. (F) Concentration series of carbenoxolone was injected over a hPANX1 coated surface for 60 seconds followed by a 90 second dissociation time. (G) The equilibrium curve does not have sufficient curvature to accurately determine the KD. All data is representative of n=3**.**

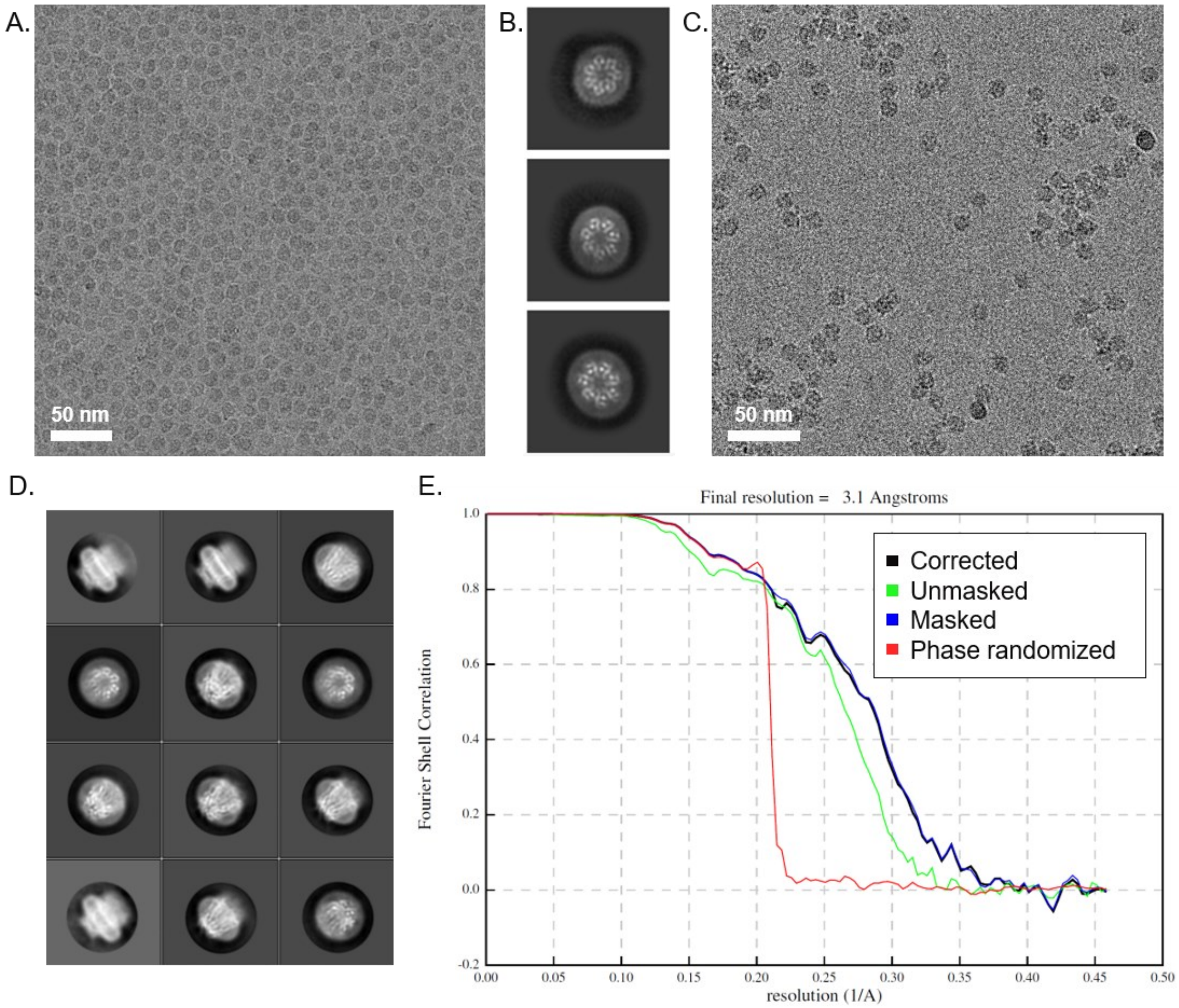

**Supplementary figure 4** - Representative micrographs, 2D class averages and Fourier shell correlation curve for the Salipro-mPANX1 datasets. (A) Representative cryo-EM micrograph and (B) 2D class averages for the dataset without fluorinated Fos-Choline 8. (C) Representative cryo-EM micrograph and (D) 2D class averages for the dataset with 0.5 mM fluorinated Fos-Choline 8. (E) Fourier shell correlation (FSC) curve plot.

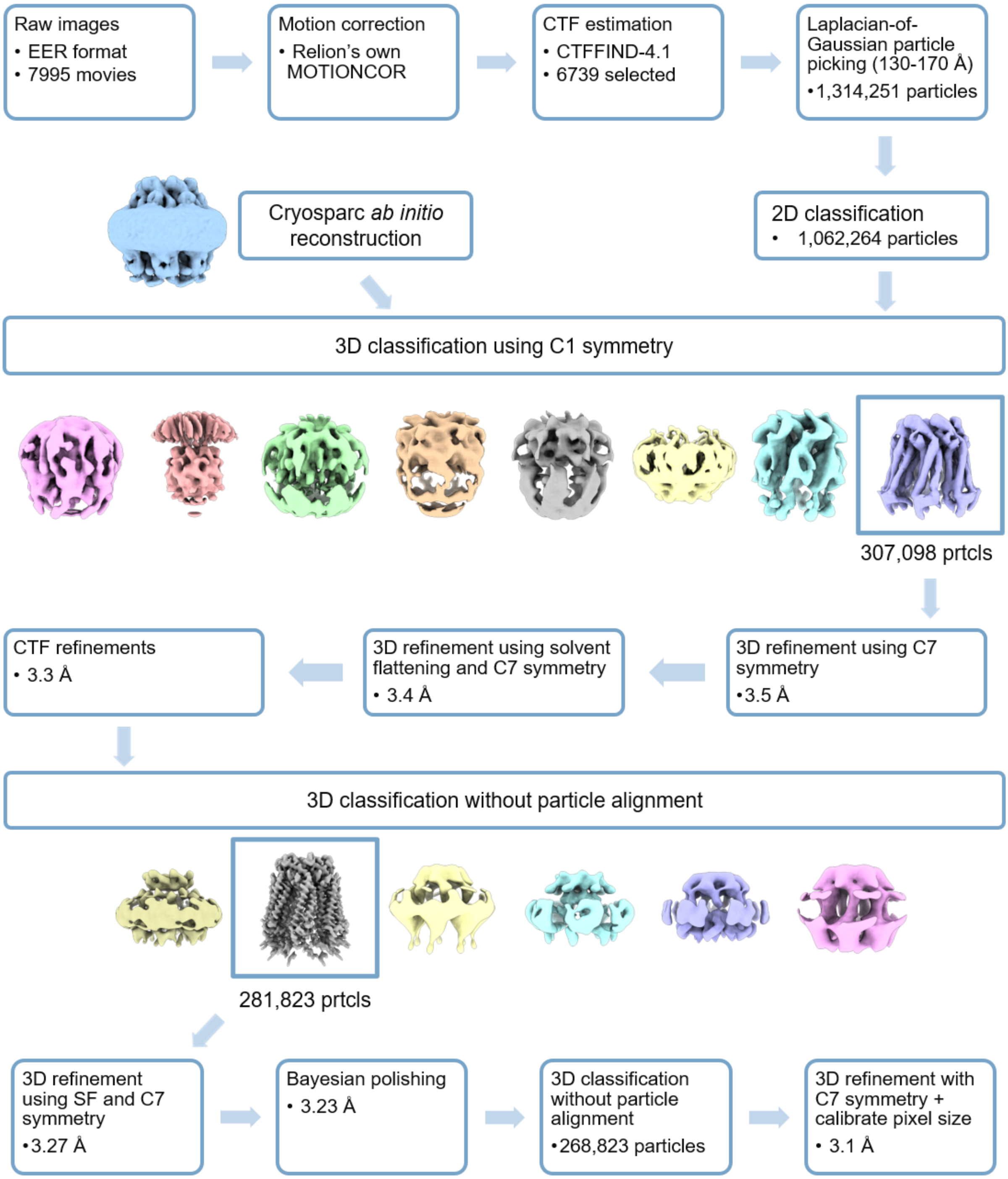

**Supplementary figure 5.** Cryo-EM image processing workflow. A detailed description of the data-analysis pipeline can be found in the Methods.

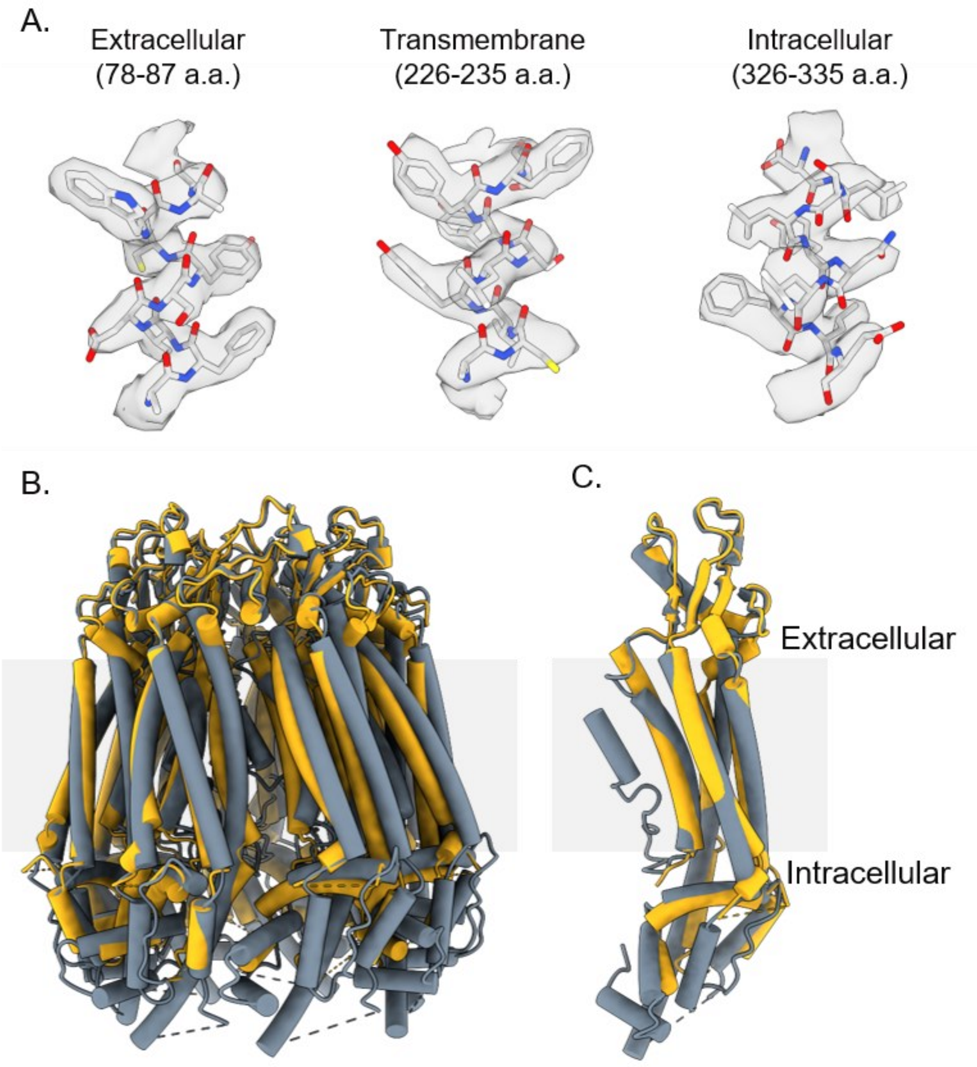

**Supplementary figure 6** – Representative density and comparison to hPANX1 (PDB code 6WBF). (A) Representative density from extracellular, transmembrane and intracellular regions. (B) and (C) Comparison of Salipro-mPANX1 (shown in yellow) heptamer (B) and protomer (C) to hPANX1 (shown in grey).

| | mPANX1 (µM) | hPANX1 (µM) |
|---|---|---|
| BzATP | 720 ± 133 | 837 ± 140 |
| Carbenoxolone | ND | ND |
| Spironolactone | 160 ± 10 | ND |

**Supplementary Table 1:** Summary of KD values determined from SPR experiments described in Figure 2B-C and Supplementary Figure 3.

| | #1 Salipro-mPANX1<br>(EMDB )<br>(PDB ) |
|---|---|
| **Data collection and processing** | |
| Magnification | 165,000x |
| Voltage (kV) | 300 |
| Electron exposure (e–/Å$^2$) | 40.24 |
| Defocus range (μm) | -0.5 to -2 |
| Pixel size (Å) | 0.727 |
| Symmetry imposed | C7 |
| Initial particle images (no.) | 1,314,251 |
| Final particle images (no.) | 268,823 |
| Map resolution (Å) | 3.1 |
| FSC threshold | 0.143 |
| Map resolution range (Å) | 2.52-5.47 |
| **Refinement** | |
| Initial model used (PDB code) | AF-Q9JIP4-F1-model_v2 |
| Model resolution (Å) | n/a |
| FSC threshold | n/a |
| Model resolution range (Å) | n/a |
| Map sharpening *B* factor (Å$^2$) | -136.316 |
| Model composition | |
| Non-hydrogen atoms | 14448 |
| Protein residues | 1792 |
| Ligands | n/a |
| *B* factors (Å$^2$) | |
| Protein | 31.948 |
| Ligand | n/a |
| R.m.s. deviations | |
| Bond lengths (Å) | 0.015 |
| Bond angles (°) | 1.507 |

| | |
|---|---|
| Validation | |
| MolProbity score | 1.36 |
| Clashscore | 6.56 |
| Poor rotamers (%) | 0 |
| Ramachandran plot | |
| Favored (%) | 97.51 |
| Allowed (%) | 2.49 |
| Disallowed (%) | 0 |

**Supplementary Table 2**. Cryo-EM data collection, refinement, and validation statistics
